# The role of the roof plate for mesencephalic trigeminal neuron guidance and position

**DOI:** 10.64898/2026.05.04.722596

**Authors:** Clemens Lumper, Artemis Koumoundourou, Max Neukum, Steffen Rauchfuss, Ulrike Kohler, Bernhard Hirt, Anthony Graham, Andrea Wizenmann

## Abstract

The mesencephalic trigeminal nucleus (MTN) contains the proprioceptive sensory neurons that innervate mechanoreceptors in the jaw closing muscles. In the chick embryo, MTN neurons are the first neurons generated in the mesencephalon. They arise bilaterally adjacent to the roof plate and then extend their axons ventrally before projecting caudally towards the rhombencephalon. MTN axons remain in a mid - dorsoventral position and pioneer the lateral longitudinal fasciculus. Notably, MTN axons never cross the roof plate, raising the question of which mechanisms underlie this restriction.

Here, we investigated the effects of tissue transplants on the guidance of MTN axons. We found that both the diencephalon and the notochord exert repulsive effects on MTN axons, which could partially explain their early trajectory. We have also analysed the potential roles of the guidance cues BMP2/4, GDF7, SLIT and NETRIN in MTN axon navigation, both *in vivo* and *in vitro*. We found no evidence for a role of BMP2/4 or GDF7 in directing MTN axons. However, SLIT-ROBO signaling was found to play a significant role. SLIT proteins are repulsive guidance cues expressed by roof and floor plate. Loss or reduced expression of ROBO2 led to aberrant axon meandering within the dorsal midbrain. Most axons eventually reoriented posteriorly, and only a small fraction crossed the roof plate. Unexpectedly, in the absence of ROBO2, MTN somata migrated into the roof plate, resulting in the loss of a defined roof plate region. Taken together, these results suggest that SLIT2-ROBO2 signaling not only prevents MTN axons from crossing the roof plate but also maintains MTN cell bodies adjacent to the roof plate. With regards to MTN neuron guidance, we conclude that additional roof plate - derived factors are likely to co-operate with SLIT proteins to prevent crossing of the roof plate. Another possibility could be that SLIT might signal through additional receptors.

## Introduction

The neurons of the mesencephalic trigeminal nucleus (MTN) are the first-born neurons of the mesencephalon (Chedotal, Pourquie et al., 1995; Hunter, Begbie et al., 2001). They are proprioceptive sensory neurons that innervate the jaw closing muscles (Corbin and Harrison, 1940; Jerge, 1963; Matesz, 1981). The innervation of the lower jaw by MTN neurons, the only primary sensory neurons in the amniote CNS (Koyama, 1987), mediates the monosynaptic jaw reflex, which triggers contractions of the muscles of mastication in response to pressure on the mandibular teeth. Hence, these neurons contribute to biting, chewing, and to a smaller extent to speech in humans (Williams, Warwick et al., 1989). The evolution of the MTN is closely linked to the evolution of the jaw (Nothcutt and Gans, 1983), and its emergence seems to be an intrinsic property of the dorsal mesencephalon (Lipovsek et al., 2017).

MTN neurons are generated bilaterally in the dorsal mesencephalon, adjacent to the dorsal midline, the roof plate (RP). Differentiation and expansion are regulated by fibroblast growth factors and WNT signaling (Hunter et al., 2001; Lipovsek et al., 2017). Their axons initially grow ventrally towards the floor plate. At the level of the mid-mesencephalon, around the sulcus limitans, they turn posteriorly towards the rhombencephalon. They cross the mid-hindbrain boundary, enter the hindbrain and exit through the trigeminal ganglion in rhombomere 2 (Hiscock, 1986). The first MTN neurons differentiate around Hamburger and Hamilton (HH; Hamburger and Hamilton, 1951) stage 13-14 and begin extending their axons ventrally at HH stage 14-15 (Chedotal et al., 1995). These first MTN axons pioneer the lateral longitudinal fascicle (LLF), one of the major axon tracts of the early axon ‘scaffold’ that forms during brain development (Chedotal et al., 1995; Easter, 1993; 1994; Wilson et al.,1990; Windle et al., 1935). However, the mechanisms that guide these pioneer axons remain only partially understood. Several axon guidance proteins have been implicated in MTN axon navigation. The initial ventral growth of MTN axons is promoted by the chemoattractant molecule Netrin, expressed in the floor plate (Tessier-Lavigne and Goodman, 1996). This ventral trajectory is further shaped by the chemo - repulsive proteins Draxin and Slit2. Draxin is expressed in a dorsal to ventral gradient in the dorsal midbrain and acts as a repellent (Naser et al., 2009). Slit2 is expressed in the roof plate in both mouse (Yuan et l., 1999) and chick (Molle et al., 2004). In chick, MTN axons avoid Slit2-expressing cells, suggesting that Slit2 contributes to preventing MTN axons from crossing the roof plate and floor plate (Molle et al., 2004).

The roles of Slit proteins and their Robo receptors in axon guidance have been studied extensively in spinal cord (see reviews: Blockus & Chedotal, 2016; Chedotal, 2019; Kaprielian et al., 2001; Blokhus et al., 2016; Guthrie, 2004; Ypsilanti et al., 2010). The floor plate serves as a source of attractive and repulsive guidance cues (Harris et al., 1996; Kennedy et al., 1994; Mitchell et al., 1996; Serafini et al., 1996; Serafini et al., 1994; Tessier-Lavigne et al., 1988; Brose et al., 1999; Colamarino & Tessier-Lavigne, 1995; Kidd et al., 1999; Stoeckli et al., 1997; Yuan et al., 1999; Zou et al., 2000). In the mouse spinal cord, Robo1 predominantly regulates the growth of ventral axon tracts and prevents midline crossing of the axons of the medial longitudinal fascicle (MLF), whereas Robo2 directs axons of more dorsally located tracts, such as the LLF (Evans et al., 2015; Kidd et al., 1998; Kim et al., 2011, Long et al., 2004; Mambetisaeva et al., 2005; Reeber et al., 2008; Sabatier et al, 2004; Simpson, Bland, et al., 2000; Simpson, Kidd, et al., 2000). In spinal commissural axons, both Robo1 and Robo2 have distinct and converging functions, while additional receptors likely contribute to Slit-mediated repulsion (Jaworiski et al, 2010). Thus, across species, including Drosophila, Slit functions as a key midline repellent (Bhat et al., 2007; Kidd et al., 1999; Rajagopalan, Nicolas, et al., 2000; Rajagopalan, Vivancos, et al., 2000; Simpson, Bland, et al., 2000; Simpson, Kidd, et al., 2000). In addition to its expression in the roof plate of midbrain, SLIT1 and SLIT2 are also expressed in the ventral midbrain and floor plate (Yuan et al., 1999, Molle et al., 2004). The floor plate has been shown to repel MTN axons (Tamada et al., 1995). Slit proteins are likely to maintain also MTN axons at a distance from the floor plate. Positioned between the roof and floor plate, MTN axons then turn posteriorly toward the hindbrain pioneering the LLF (Brose et al., 1999; Chen et al., 2001; Hu, 2001; Li et al., 1999; Nguyen Ba-Charvet et al., 1999; Simpson, Kidd, et al., 2000; Shu and Richards, 2001; Wu et al., 1999). This posterior turn may be also influenced by strong Slit expression in ventral diencephalon (Molle et al., 2004. Other roof plate-derived signals may also contribute to this process. BMP4, BMP7 and GDF7 are also expressed in the roof plate, and BMP4 and BMP7 have been shown to repel commissural axons in the spinal cord (Augsburger, 1999, Butler, Dodd, 2003). In addition to their effects on axons, Slit proteins also regulate neuronal positioning. They prevent for example, motor neuron cell bodies from crossing the floor plate in the spinal cord (Kim et al, 2011; 2015). Thus, Slit2 may have a dual role in dorsal midbrain: repelling MTN axons from the roof plate and preventing MTN cell bodies from migrating into it, thereby maintaining the bilateral position of MTN neurons and axons.

Here, we investigated the role of roof plate-derived SLIT2 and BMPs in preventing MTN axons from crossing the midline and in maintaining the proper positioning of MTN neuronal cell bodies within the mesencephalon. Using a combination of *in vivo* and *in vitro* assays, together with a ROBO2 receptor knockdown, we examined the influence of different brain regions, signaling centers, and guidance cues on MTN axon pathfinding and neuronal positioning.

Our results confirm that the roof plate, notochord, floor plate and diencephalon act as repulsive sources for MTN axons and that Slit2 contributes to this repulsion, although it is not the only factor preventing roof plate crossing. Furthermore, we identify an additional role for SLIT2 in constraining MTN neuronal somata to positions lateral to the roof plate.

## Methods Transplantations

Fertilized chicken eggs (Lohmann Brown) were purchased from Artländer Geflügelhof, Quakenbrück, Germany. They were incubated at 38°C with 65% humidity until the desired developmental stage after Hamburger-Hamilton was reached (Hamburger & Hamilton, 1992). The embryos were prepared as described in Huber et al. (Huber et al., 2013). In short, embryos were incubated up to Hamburger and Hamilton stage (HH) 13-14 (Hamburger and Hamilton, 1951), then a small hole was cut into the eggshell. The vitelline membrane above the embryo was opened and a small whole was cut into the developing medial mesencephalon, where MTN axons pass using sharpened tungsten needles. Cell aggregates or prepared donor tissue that was labelled with Orange Cell Tracker (Molecular Probes) was maneuvered into place with tungsten needles. The hole in the eggshell was closed with flexible tape and incubated until HH 16 - 18.

### Cell aggregates

Cells from the 293-T line (human embryonic kidney cells), the Ton-293-Netrin-line, and Xenopus Slit expressing EBNA cells (human embryonic kidney cells from the HeEK293 line; xSLit, Wu et a., 1999) were cultured in DMEM with 20% FCS and 1% Pen/Strep. After 3 to 4 days, when cells covered the entire ground of the petri dish, they were treated with trypsin / EDTA (8 minutes), collected, washed and labelled with orange Cell Tracker. They were diluted to 10^3^ cells / ml and drops of 20 ml were brought onto a lid of a 6 cm petri dish that was filled with 2 ml PBS. The lid was inverted over the dish and the cells were incubated for 20 h at 37°C at 5% CO_2_ and 50% relative humidity prior to transplantation or coculturing. The cells formed aggregates within the drops and were used for transplantation (Kennedy et al., 1994).

### Cell Tracker Labelling of living cells

To label cells and chicken tissue, orange Cell Tracker (Molecular Probes) was employed. 10 mM of Cell Tracker was dissolved in DMEM and the cells and tissues were incubated for 20 min at 37 °C. They were washed 3 times in DMEM.

### Cocultures

To analyse the growth behaviour of MTN axons were cultured opposite different types of tissue or cell aggregates from different cell lines in collagen. MTN explants consistent always of a roof plate (RP) in the middle of the explant and MTN neurons to both sides of the RP. Hence, the side of the explant not encountering foreign tissue could be used as a control side where MTN axons grew without an obstacle on their way.

### In vivo transplantations

Various types of tissues and factor expressing cell aggregates were grafted into the dorsal midbrain at HH stage 13 - 14 to test how MTN neurons react *in vivo* (see Li et al., 2005). In short, fertile eggs were incubated at 37oC to the required HH stage. The egg was windowed and the embryo visibility was enhanced by sub-blastodermal injection of diluted India Ink (1:20 in phosphate buffered saline). The viteline membrane was opened and tissue from the midbrain half receiving the transplant was cut out in the size of the transplant. Donor pieces from different brain regions, cell aggregates or beads (see below) were with an orange Cell Tracker dye (CMTMR, Molecular Probes) before transplantation into dorsal midbrain. Grafts were oriented into the cut with tungsten needles. The egg was sealed, incubated to the required stage, and the embryo was removed and fixed for further treatment. Midbrains where the transplant had not intergrated well were discarded.

### BMP2/4 beads

To test the effect of BMP4 on the growth direction of MTN axons heparin coated beads carrying BMP2 were transplanted into their pathway. BMP4 and BMP2 are related to the Drosophila decapentalplegic subfamily and have been shown to antagonize FGF function (Neubüser et al., 1997; Niswander and Martin, 1993). Thus, they have similar effects.

### *In ovo* electroporation

Fertilized chicken eggs were incubated at 38 °C in a hatching chamber in a humidified atmosphere. Embryonic stages were determined according to Hamburger and Hamilton (Hamburger and Hamilton, 1951). At HH 12 - 14, 2 ml of egg albumen were removed, and a window was opened in the eggshell to expose the living embryo. si-ROBO2 expression constructs were injected into the chick neural tube at HH 12 - 14. The si-ROBO2 (1 µg / µl) constructs were mixed with a GFP or a VenusLyn (kind gift of Dr. H. Lieckert, Münich) reporter plasmid in a 1 : 0.9 ratio. The electroporation was performed as described by Itasaki et al., Momose et al. and Huber et al. (Huber et al., 2013; Itasaki et al., 1999; Momose et al., 1999). Briefly, the cathode was placed on the ventral side of the neural tube and the anode on the dorsal side of the mesencephalon. Four 10 ms / 15 V pulses were applied to electroporate dorsal midbrain cells that will develop into MTN neurones. Approximately, 500 ml phosphate buffered saline (PBS) was added. Eggs were sealed and incubated until they reached HH stage 15 - 18. Then, the embryos were dissected out and fixed in 4% PFA (in PBS). The unelectroporated midbrain half served as internal control. Electroporations with only GFP or VenusLyn did not have an effect on axonal growth (data not shown, see also Moschou et al.; 2026).

### siROBO2 Constructs

Three siROBO2 constructs were subcloned into the pSilencer1.0 vector, following the manufacturers (Ambion) instructions. The target sequences (5’-GCGTCTCAATACCATTTATCT - 3’, 5’-GGAAATCACAACAGCACAAGC-3’ and 5’-GTGGTGCATGTTGGGTAATT-3’) were selected using http://www.rnaiweb.com (now defunct). All three vectors were electroporated simultaneously with an additional vector expressing GFP as a means to report successful transfection.

### Immunohistochemistry

For whole-mount immunostainings, embryos were fixed overnight in 4% PFA, washed five times 30 min at 4 °C in PBS plus 0.1% TritonX-100 and 10% FCS and then incubated for 2-4 days at 4 °C in primary antibody solution. Then, after five 20 min washes, embryos were incubated with fluorochrome-conjugated secondary antibodies for up to 2 nights at 4 °C. Embryos were washed again five times, shortly fixed in PFA and then flat-mounted. Primary antibodies used were 3A10 a neurofilament associated antibody (The 3A10 antibody developed by Jessell, T.M., Dodd, J. and Brenner-Morton, S. was obtained from the Developmental Studies Hybridoma Bank, created by the NICHD of the NIH and maintained at The University of Iowa, Department of Biology, Iowa City, IA 52242)) and anti-GFP (Mobitech). Secondary antibodies were anti-mouse and anti-rabbit Cy2 and Cy3, (Jackson ImmunoResearch Laboratories. Inc. 115-165-003, 111-226-003).

### Whole-mount mRNA *in situ* hybridization

Whole-mount *in situ* hybridization was performed according to Henrique et al. (Henrique et al., 1995). In short, an antisense mRNA sequence of ROBO2 was synthesized *in vitro* using the T7 RNA polymerase and digoxigenin (DIG) coupled UTPs in an RNAse-free buffer solution (plasmid with ggROBO-2 was a kind gift of Avihu Klar). The fixed tissue was dehydrated, rehydrated and treated with phosphate buffered saline containing 6% hydrogen peroxidase, then washed in the hybridisation buffer and incubated over night at 70oC in hybridisation buffer with the DIG coupled anti sense ROBO2 constructs. Then, the tissues were washed and incubated with an alkaline phosphatase conjugated anti DIG antibody (Roche Diagnostics). NBT and BCIP (Roche Diagnostics) were used as substrates for the staining reaction.

### Flat-mounts

For flat mounts of the midbrain at HH14 - 18, the midbrain was separated from the rest of the embryo. Then the mesenchyme surrounding the midbrain was removed. A cut along the roof plate allowed the flattening of the tissue on a slide with the basal side facing the cover slip. The tissue was embedded in PBS and glycerol (50% each with 0.01% Triton). This mixture led to a passive ‘clearing’ of the tissue (Neckel et al., 2016). Thus, labelled cells and axons could be much better identified.

### Growth analysis and Statistics

The growth of the axons was classified in three groups. Axons growing more or less radially away from their source (defined by us as not turning more than 20^0^ towards or away of the tested tissue) were classified as normal and categorized as group I. Axons taking a turn of more than 20^0^ towards the tested tissue were categorized as attracted and put into group II. Group III consisted of axons repelled by the tissue they were confronted with, resulting in the axons having turned more than 20^0^ away from the tissue. This classification of the axons was performed at control - and experimental side. The length of the axons was measured with a rotatable scale in an ocular.

All statistical analysis was done using IBM SPSS Statistic 29.0.2.0 software. To test for significance of MTN axon outgrowth direction when confronted with different tissues or cell lines the non-parametric Mann - Whitney U test (also called the Mann – Whitney - Wilcoxon rank test) was used for both co - cultures and transplants. Except for the roof plate co-cultures, for which a Wilcoxon Signed Rank test was used to test for significance of the axon outgrowth direction. The roof plate co-cultures were the only samples where counts of axons for the exposed and control side could be matched.

## Results

### Diencephalic and Notochord Tissue Repel MTN Axons

The development of MTN neurons at different HH stages is shown in Figure 1. The first neurons differentiate at HH13. At HH14, axons begin to grow ventrally, away from the roof plate (Figure 1A, arrows). At HH15, the first axons reach the sulcus limitans and turn caudally toward the rhombencephalon (Figure 1B). These axons pioneer the longitudinal lateral fascicle (LLF; Hiscock et al., 1986; Chedotal et al., 1995). At HH16, the position of the LLF at the level of the sulcus limitans becomes recognisable (Figure 1C), while the medial longitudinal fascicle (MLF) has already formed. By HH 17 (Figure 1D), the LLF is well defined. At all stages, MTN axons remain confined to specific tracts: they do not cross to the contralateral half of the midbrain, neither ventrally nor dorsally, nor do they enter the diencephalon.

**Figure 1:**
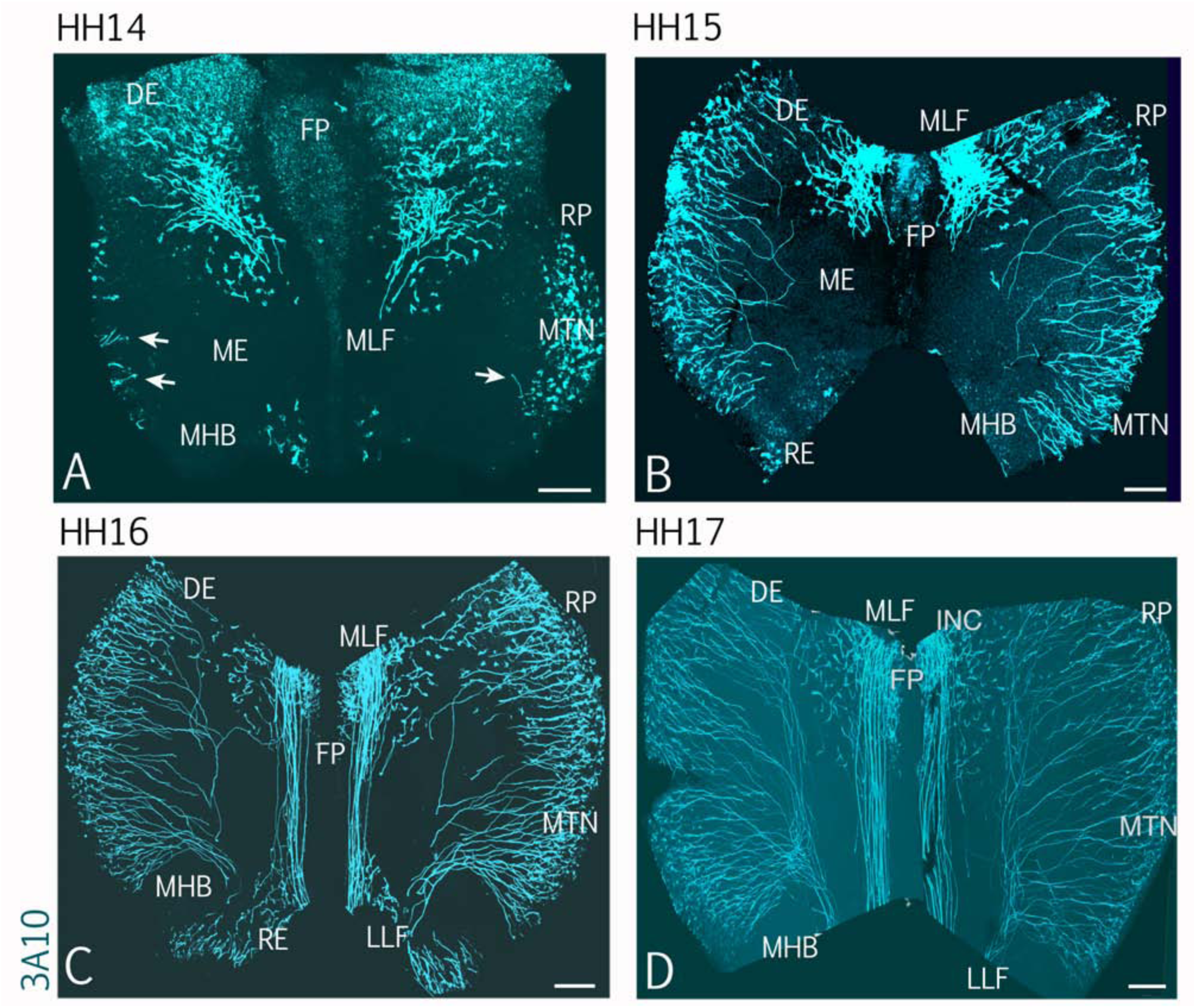
Pathway of mesencephalic trigeminal axons at different HH stages. A-D: Midbrains of different HH stages. The midbrains were cut along the roof plate and flat mounted. Neurons and axons were labelled with green immunostaining (3A10). A. At HH 14 the first MTN neurons differentiate left and right of the roof plate. Several axons growing ventrally can be seen (arrowheads). B. At HH15 some MTN axons have reached the medial midbrain and turned posteriorly towards the rhombencephalon. C and D: The LLF formed by MTN neurons becomes visible at HH16 (C) and can be clearly seen at HH17 (D). The MLF left and right of the FP is generated by axons from the diencephalon and is already clearly visible at HH16 (C). Scale bars: 100 µm. Abbr.: Di – diencephalon, FP – floor plate, INC – interstitial nucleus of Cajal, LLF – lateral longitudinal tract, ME – mesencephalon, MLF – medial longitudinal tract, MTN – mesencephalic trigeminal nucleus, RE – rhombencephalon, RP – roof plate.

To investigate the influence of the diencephalon and notochord on MTN axon growth, we performed transplant and co-culture experiments. For co-cultures, dorsal midbrain tissue containing MTN neurons was cultured with dorsal midbrain, diencephalon or notochord tissue (all from HH 10-12 embryos).

*In vitro*, 67% of MTN axons growing from dorsal midbrain explants turned away from adjacent diencephalon tissue (n = 13; SD 14%, p<0.001, Figure 2A, I). In contrast, axons on the opposite (control) side, not exposed to diencephalic tissue, grew predominantly straight, with only 21% showing a turning behaviour consistent with repulsion. The repulsion effect of the diencephalon was confirmed in vivo. Transplantation of diencephalic tissue into the path of MTN axons in embryos resulted in 77% of axons being repelled during their ventral growth towards the floor plate (n = 4, SD 3%, p<0.006,Figure 2C, J,). Transplanted notochord tissue had a similar effect (Figure 2E, J) with 84% MTN axons being repelled (SD: 16%, p<0.056, n = 2). Remarkably, MTN axons consistently grew around the notochord transplant in a wide circle rather than simply avoiding direct contact. As the notochord is known to induce floor plate tissue (Roelink et al., 1994; Placzek et al., 2000; Patten and Placzek, 2002), it is likely that the surrounding tissue adopted floor plate-like properties and expressed guidance cues such as Slit and Netrin. Our results suggest that repulsive cues, most likely Slit proteins, dominated over any potential attractive effects of Netrin (not directly tested). Together, these results indicated that both diencephalic and notochord tissue exert repulsive effects on MTN axon growth *in vitro* and *in vivo*. We next tested the effect of Netrin - expressing cells in co-cultures and *in vivo* with transplantation assays (Figure 2B, D, I, J). *In vitro,* 67% of MTN axons were attracted toward Netrin1 - expressing HEK293 cells (n = 11). On the control side, most axons extended radially away from the MTN explant (Figure 2B), although some fibres appeared to turn toward the Netrin source, possibly due to diffusion of Netrin across the culture. Quantitative comparison confirmed that attraction toward Netrin - expressing cells was significantly greater than in controls (Figure 2I; SD 17%, p<0.001).

**Figure 2:**
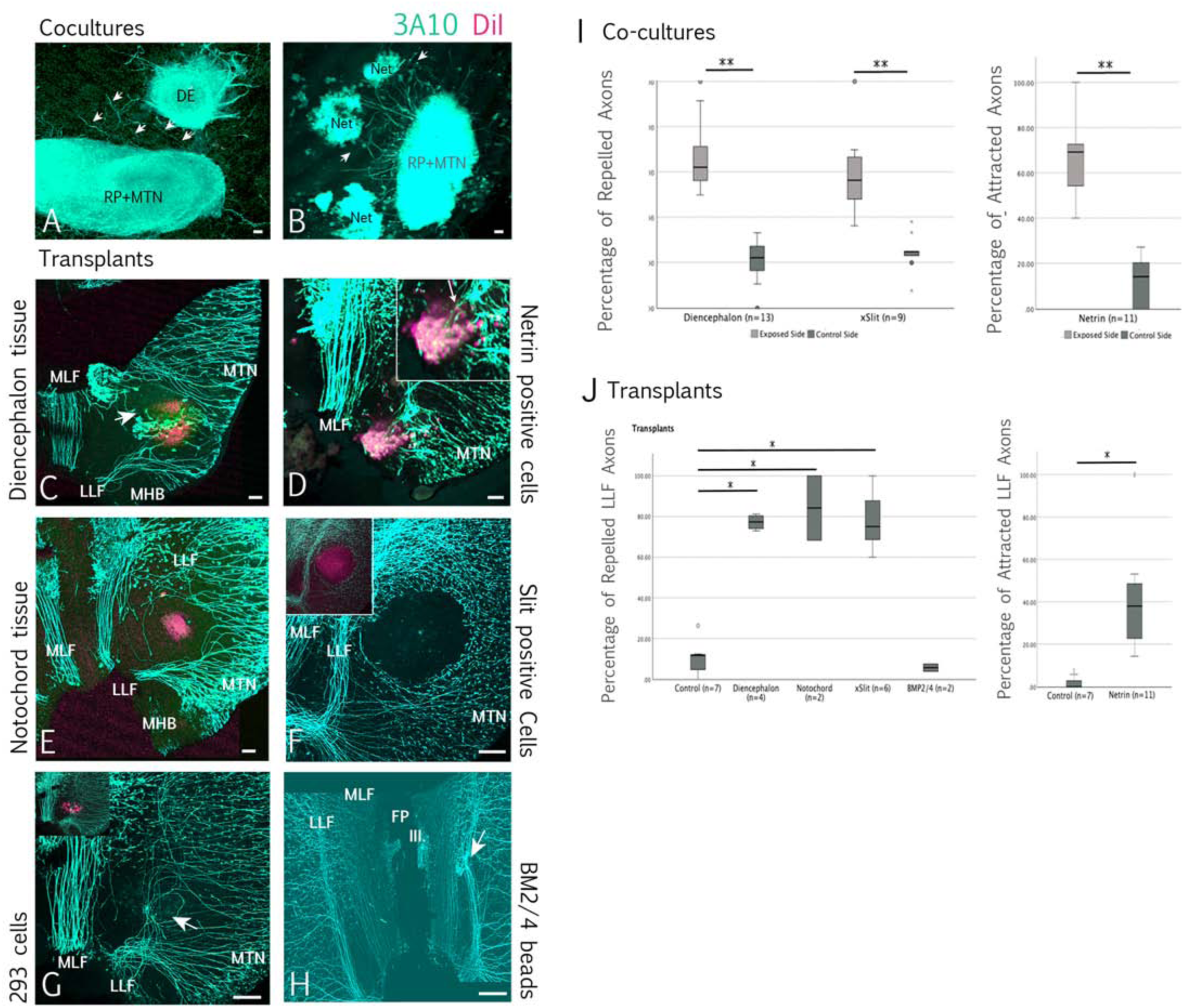
Influence on MTN axon growth by different regions/tissue?? and proteins. A, B: Cocultures of dorsal midbrain containing MTN neurons with diencephalon (A) and (B) with Netrin expressing cells. C - H: Transplants of different tissues, cell aggregates and beads into dorsal midbrain. Transplanted tissue and cells are labelled red (DiI), neurons and axons are green (3A10). A: MTN axons confronted with diencephalic tissue avoided diencephalic explants in 67% of all examples (I). B: Netrin expressing cells had mostly an attractive effect on MTN axons (67%, I). C: Diencephalic transplants showed a strong effect on the growth direction of MTN axons. The axons avoided the transplant (arrow; 77 %; I). D: Cell aggregates expressing Netrin did not disturb MTN axons path much when transplanted into their way (see picture magnification at the top right). The axons crossed the cell aggregate. 44% of the axons were attracted towards Netrin expressing cells (J). E: Transplants of notochord tissue were avoided by MTN axons in an area wider than the transplant (compare size of red cells and axon growth). In average 84% of MTN axons avoided the transplant (J). F: Cell aggregates expressing Slit2 proteins also had a rejecting effect on the growth of MTN axons in 78% (J). Here, the axons grew close around the cell aggregate expressing Slit2. The size of the transplant is indicated at the top left. Slit2 expressing cells are labelled in red. G: Cell aggregates consisting of 293 cells (red) as control did in some cases cause a slight axon growth irritation (arrow; 10%, J) but all axons crossed the transplant and formed the LLF. The transplant location is shown at the top left. H: Beads loaded with BMP2/4 transplanted into midbrain seemed to attract some MTN axons (arrow) but did not misdirect them. The axons grew posteriorly and formed the LLF. I, J.: Statistical analysis of the outgrowth behaviour of MTN neurons encountering different tissue or cell aggregates. I: In Cocultures the repellent effect of diencephalic tissue or Slit expressing cells is dominant. Cells expressing Netrin have an attractive effect on MTN axonal growth. J: The statistical evaluation of transplants of notochord, diencephalon or xSlit expressing cells showed that all three had a strong repellent effect on MTN axon growth. Beads expressing BMP beads did have no repellent effect comparable to the control tissue. Netrin expressing cells did attract almost half of the MTN axons. Scale bars: 100 µm. Abbr.: III. – nervus oculomotorius, DE – diencephalon, FP – floor plate, LLF – lateral longitudinal tract, MLF – medial longitudinal tract, MHB – mid - hindbrain boundary, MTN – mesencephalic trigeminal nucleus, Net - Netrin expressing cells, RP – roof plate.

*In vivo* transplantation of Netrin - expressing cells into the MTN axon pathway produced similar results (Figure 2D). 45% of axons were attracted towards the transplant (n = 11, SD 29%, p<0.001, Figure 2J). As control, aggregates of non - transfected HEK293 cells were transplanted into dorsal midbrain (Figure 2G). While these occasionally caused minor disturbances on axon trajectory, the LLF was formed normally, and only 10% of axons exhibited repulsion (n = 7, SD 8%, Figure 2J).

Previous studies demonstrated that Slit proteins act as strong repellents for MTN axons. Molle et al. (2004) showed that 74% of the MTN axons avoided Slit - expressing cells in co-culture assays and Farmer et al. (2008) reported that in Slit1/ 2 or Robo1 /2 double mutants, the LLF loses its fasciculation and lateral positioning, with axons displaying aberrant, meandering trajectories. To confirm the repulsive effect of Slit proteins, we transplanted Slit - expressing cells into the pathway of MTN axons. MTN axons consistently avoided these cells and instead grew around them (Figure 2F). Quantitative analysis showed that 78% of MTN axons encountering Slit - expressing cells were repelled (Figure 2J; n = 6, SD13%, p<0.001), confirming their high sensitivity to Slit - mediated repulsion. Taken together, these findings demonstrated that diencephalic tissue, notochord and Slit - expressing cells exert strong repulsive effect on MTN axon growth.

### Influence of the roof plate on growth direction of MTN axons

To investigate the influence of the roof plate on the MTN axon direction, roof plate tissue from HH10-12 embryos was isolated and either used in co-culture assays or transplanted into dorsal midbrain of embryos at approximately HH10-11. The co-culture experiments showed that the roof plate exerts a repulsive effect on the MTN axon (Figure 3A;). 79% of MTN axons (n = 18, SD 18%, p<0.001, Fig.3C) extending from explants facing the roof plate tissue turning away (arrows in Figure 3A). In contrast, axons at the opposite (control) side that were not exposed to roof plate tissue, extended radially away from the explant (Figure 3A, arrowheads). Consistent with the *in vitro* findings, *in vivo* transplantation of roof plate tissue into the dorsal midbrain resulted in a clear repulsive effect on MTN axons (Figure 3B). Quantitative analysis showed that 80% of host MTN axons were repelled by the transplant (n = 8, SD 18%, p<0.001, Figure 3D). MTN axons avoided the transplanted roof plate tissue and the axons located anterior to the graft altered their trajectory dramatically (Fig 3B). In some cases they even grew into the diencephalon (see arrow in Fig. 3B). This repulsive effect was consistently observed. Together, these results demonstrate that the roof plate acts as a strong repulsive cue for MTN axons and significantly influences their growth direction.

**Figure 3:**
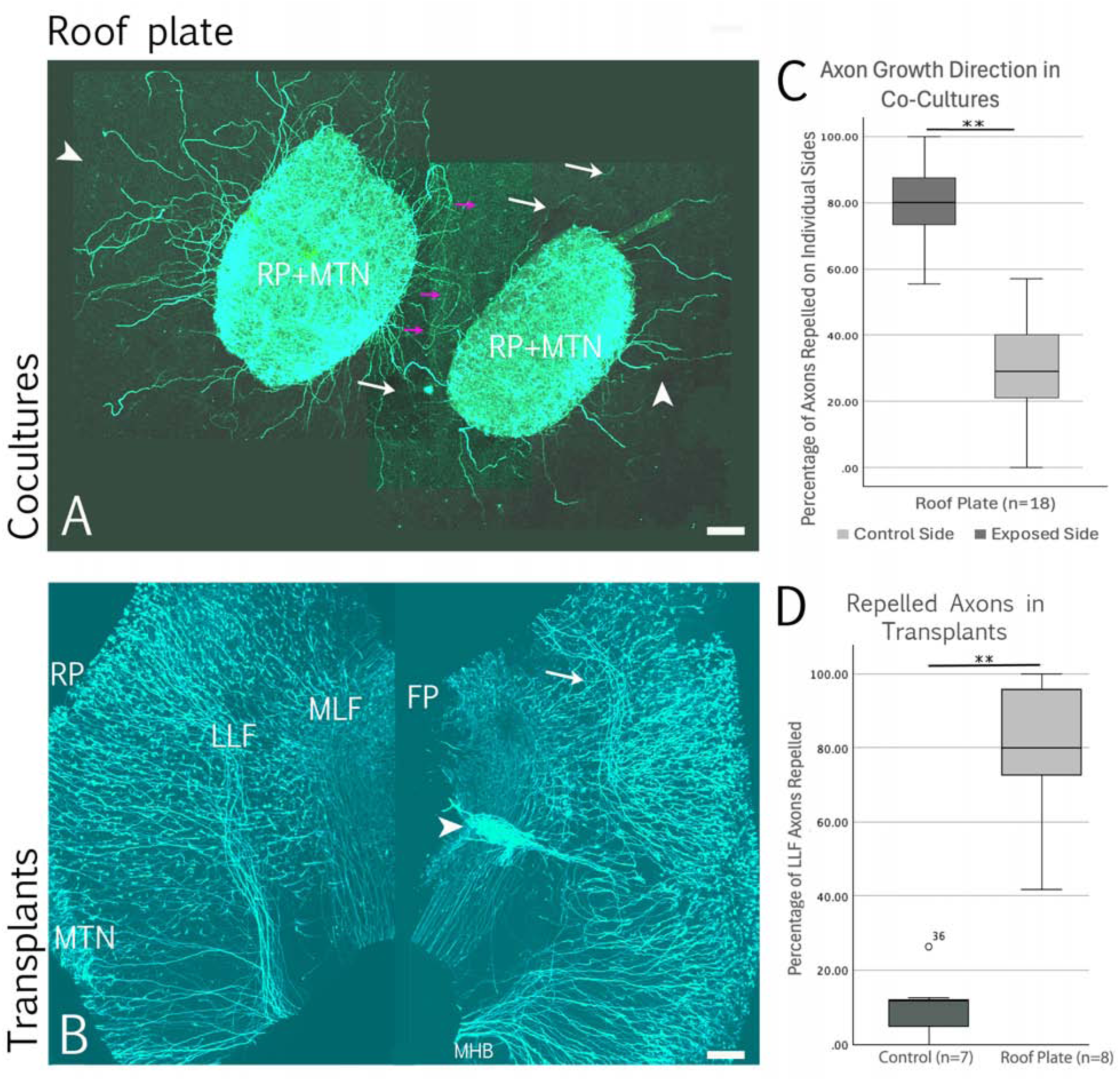
Effect of roof plate cocultures and transplants on MTN axonal growth. A: Midbrain roof plate tissues containing MTN neurons were co-cultured. The MTN axons that grew out of the explants were repelled by the opposite dorsal midbrain explant (see white and pink arrows). MTN axons not confronted with roof plate tissue did grew more or less straight away from the transplant (white arrowheads). Statistical evaluation (C) showed that 90% of MTN axons avoided the opposite explant. B: Transplants of midbrain roof plate into the path of MTN axons resulted in a strong repellent effect on MTN axons. The axons even changed direction and grew into the diencephalon (arrow) where they still formed a LLF, D: Statistical analysis showed that in 80% the axons were repelled from roof plate transplants. Scale bars: 100 µm. Abbr.: FP – floor plate, LLF – lateral longitudinal tract, MHB – mid - hindbrain boundary, MLF – medial longitudinal tract, MTN – mesencephalic trigeminal nucleus, RP – roof plate.

### Roof plate proteins that influence axon growth direction

In spinal cord, BMP4, BMP7 and GDF7, which are expressed in the roof plate, have been shown to repel commissural axons (Augsburger et al., 1999; Butler and Dodd, 2003; review Zhao, 2003). These proteins are also expressed in the midbrain roof plate and within the dorsal region (Bobak et al., 2009; Bothe et al., 2011; Chizikov et al., 2004; GEISHA database). Other candidate repellants for MTN axons include Slit and Draxin, both of which are axon guidance molecules expressed in the roof plate and the adjacent dorsal midbrain (Islam et al., 2009; Molle et al., 2008; Naser et al., 2009; Yuan et al., 1999). To determine which roof plate – derived factors are responsible for the repelling of MTN axons we investigated the role of SLIT2, BMP4, and GDF7.

### BMP signaling has no influence on MTN axon pathway

To investigate the influence of BMP signaling on MTN axon guidance, we applied BMP2-coated beads into the dorsal midbrain. In some cases, MTN axons appeared to show a weak attraction towards the beads (Figure 2H, see arrow). However, comparison with controls (n = 7) and statistical analysis revealed no significant attractive or repulsive effect (n = 2, SD 2%, p<0.333,Figure 2 J).

Furthermore, ectopic expression of BMP4 and GDF in dorsal midbrain had no detectable effect on MTN axon growth (data not shown, see Moschou et al, 2026). As control, transplants of 293-T cell aggregates were placed into the dorsal midbrain (Figure 2G). Despite the presence of these grafts, the majority of MTN axons (88%; n = 7; SD 8) followed the normal trajectory and extended through the transplants (Figure 2J). Only a small and statistically insignificant fraction of axons appeared attracted (2%) or repelled (10%; Figure 2J). These findings indicate that BMP signaling does not play a significant role in directing MTN axons away from the roof plate.

### SLIT – ROBO Signaling Influence on MTN development

To determine whether SLIT2 signaling is required to prevent MTN axons from crossing the roof plate, we examined the expression of its receptor ROBO2. ROBO2 is expressed in chick MTN neurons (Figure 4A, B), with the earliest expression at HH15 in newly differentiated MTN cell bodies (Ratie et al., 2013). At HH18, the mesencephalic roof plate is clearly visible between the bilateral MTN populations (Figure 4A). This separation is also evident at earlier stages (HH15 - HH18; arrowheads in Figure 5 A-C, n = 16). In wildtype embryos, MTN axons grow away from the roof plate, while the cell bodies remain positioned on either side.

**Figure 4:**
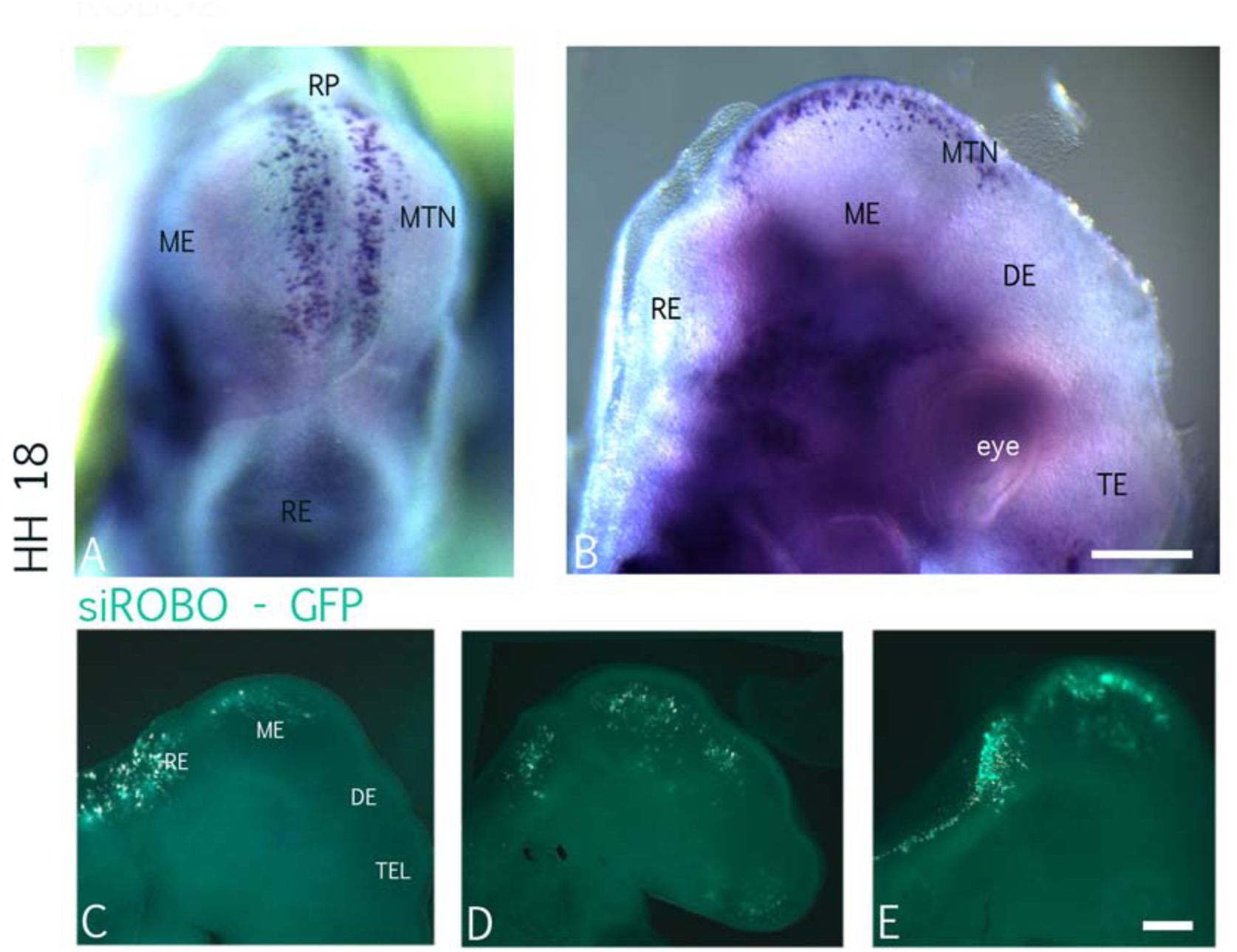
*ROBO2* mRNA expression in MTN neurons. A: View onto the top of an HH18 mesencephalon. *ROBO2* mRNA (blue) is clearly present in MTN neurons left and right of the mesencephalic roof plate. B: Side view of an HH18 head. No other cells than MTN neurons express *ROBO2* in the dorsal midbrain or other brain areas at that developmental stage. C - D: Images of chick heads at HH 16/17 successfully electroporated with *siROBO2* + GFP. In A and C MTN neurons are GFP positive. In B, GFP positive cells are too lateral of the roof plate to be present in MTN neurons. Scale bar: 100 µm. Abbr.: DE – diencephalon, ME – mesencephalon, MTN – mesencephalic trigeminal nucleus, RE– rhombencephalon, RP – roof plate, TE – telencephalon.

**Figure 5:**
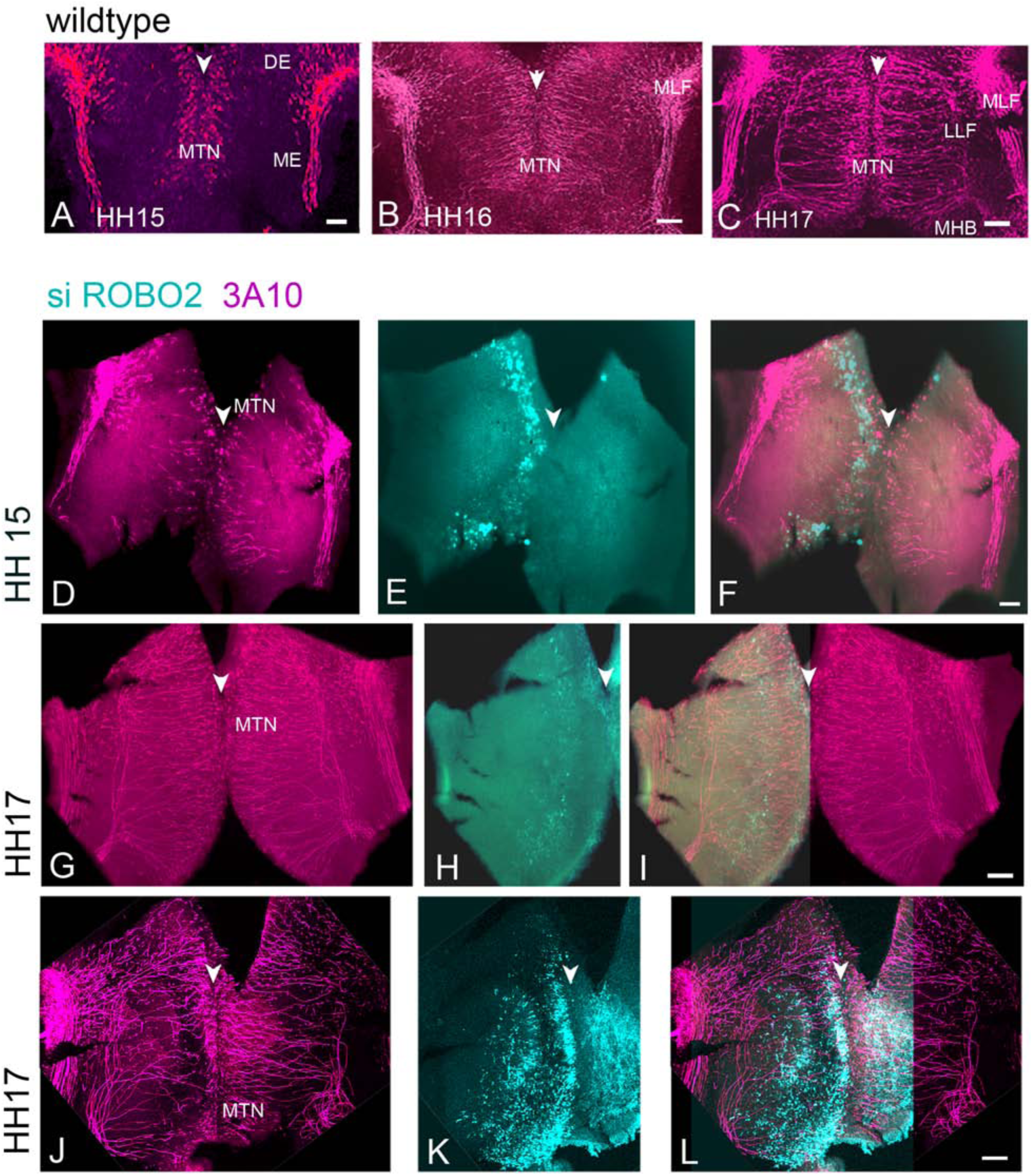
Effect of *siROBO2* expression on MTN axonal growth direction. A - C: Wild type axonal growth at HH15, HH16, and HH17 in midbrain. The arrowhead points to the roof plate clearly visible between the MTN neurons left and right of it. Midbrains were cut at the floor plate and flat mounted, neurons were labeled with 3A10 (red). D - L show examples of *siROBO2* expression in dorsal midbrain cells in different distances to the roof plate. D, G, and J show the course of MTN axon, E, H, and K the *siROBO2 –* GFP expression, and F, I, and L the overlay of axons and GFP expressing cells. D - F: *siROBO2* expressing cells at HH 16 lateral to the roof plate did not disturb the course of MTN axons. Only few to none of the MTN neurons expressed *siROBO2* GFP (F). G - I: Cells expressing s*iROBO2* in lateral midbrain at HH17 do not disturb MTN axon course (H, I). Both LLFs formed and crossed the mid-hindbrain boundary (G, I). The GFP -*siROBO2* expressing cells are ventrally of the MTN location (H). J - L: Another example of *siROBO2* transfected cells in dorsal midbrain at HH17. This example shows GFP positive cells near the roof plate (K, arrowhead) and thus several MTN neurons should contain *siROBO2-*GFP (L). The MTN axon paths in control and transfected halves are very similar. LLFs in both midbrain halves were formed. Scale bars: 100 µm. Abbr.: DE – diencephalon, LLF – lateral longitudinal tract, ME – mesencephalon, MHB – mid - hindbrain boundary, MLF – medial longitudinal tract, MTN – mesencephalic trigeminal nucleus, RP – roof plate.

To investigate the functional role ROBO2, we designed three siRNA constructs targeting ROBO2 and electroporated them into dorsal midbrain cells adjacent to the roof plate (4C – E). MTN axon trajectories were visualized in flat - mounted midbrains using 3A10 immunostaining.

Transfection efficiency varied between experiments (compare Figure 5D, and Figure 6). In cases where transfection was restricted to cells ventral or lateral to the MTN population (Figure 5E, H, K, n = 9), MTN axons followed their normal trajectory and the roof plate remained clearly defined (Figure 5D, G, J; arrowheads). In contrast, when MTN neurons were transfected with siROBO2, LLF formation was disrupted in 10 out of 11 midbrain halves (Figure 6A, E, H, K, L; arrows). Axon growth appeared disorganized. The transfected axons no longer extended directly toward the floor plate, and many failed to turn caudally (Figure 6A, E, H, K). Some axons grew aberrantly along the roof plate (Figure 6M), and a small portion crossed the midline (5/11; Figure 6M). These defects were most pronounced when both, roof plate and adjacent MTN neurons were targeted by transfection (Figure 7B, F, I, L). In most cases, only one side of the midbrain was transfected, allowing the contralateral side to serve as an internal control, which consistently exhibited normal LLF development (Figure 6A, E, H, K, L).

**Figure 6:**
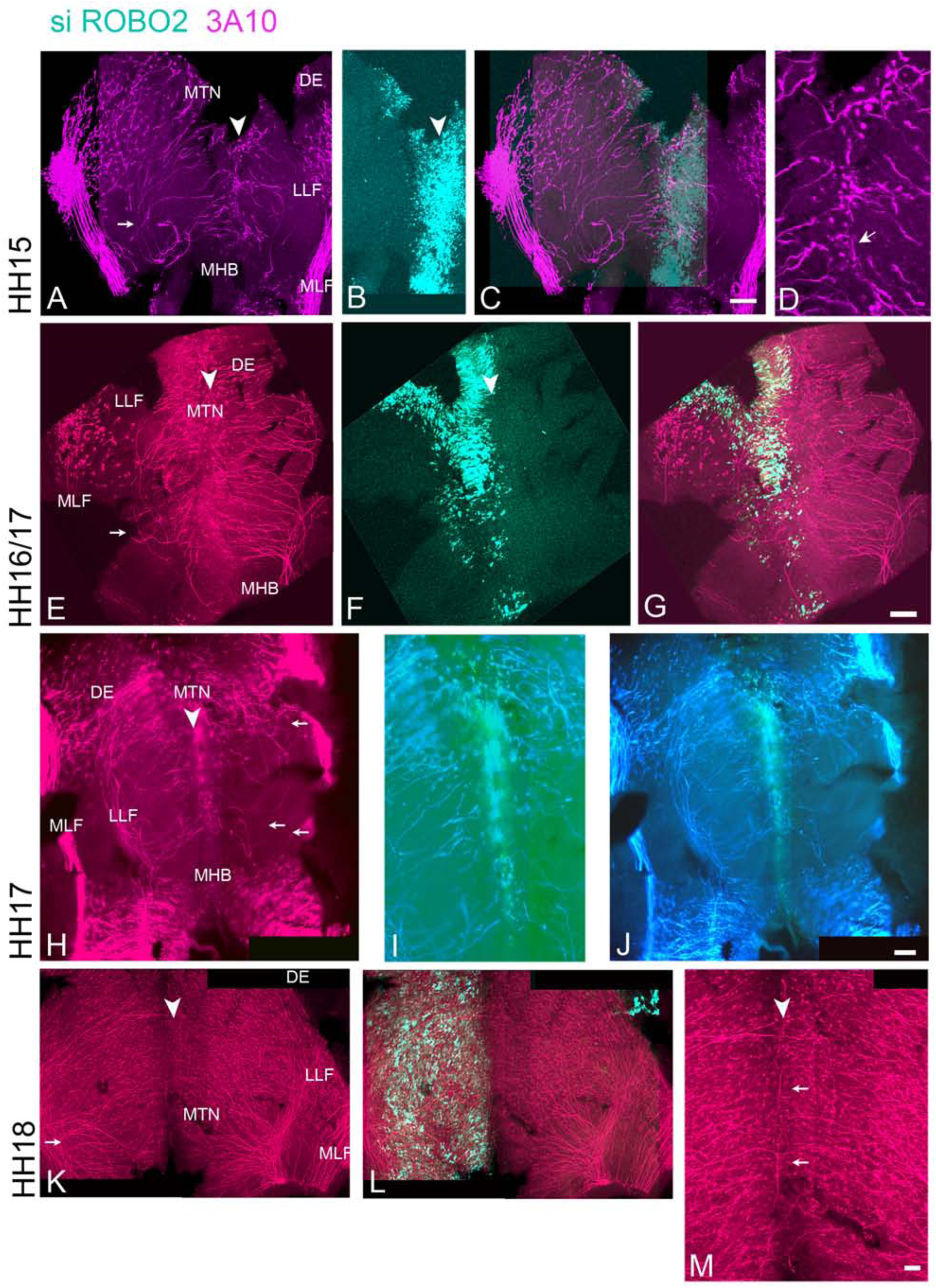
Effect of *siROBO2* expression on MTN neurons. A - M show examples of *siROBO2* transfection in MTN neurons and roof plate (A - J) or only MTN neurons (K - M) that influenced axonal growth and MTN cell location. Arrowheads in A, E, H, and K point to the location of the roof plate. In A, E, and H the straight growth of MTN axons towards the medial midbrain was disturbed within the midbrain half that was transfected with *siROBO2-GFP*. MTN axons in the transfected midbrain half seem to have difficulties to form a proper LLF compared to the non - transfected half. Some axons even grew anteriorly (arrows in E and H). K - M show an example where *siROBO2* transfected MTN axons grow almost normally and form the LLF. In A, E, H, and K the roof plate is not as clearly visible as in control midbrains and some axons (arrow) grow within it (M, D). Scale bars: 100 µm. Abbr.: DE – diencephalon, FP – floor plate, LLF – lateral longitudinal tract, MHB – mid - hindbrain boundary, MLF – medial longitudinal tract, MTN – mesencephalic trigeminal nucleus.

To assess whether ROBO2 also affects MTN neuronal positioning, we examined the organization of the roof plate. In all cases where MTN neurons were transfected (10/11), the roof plate appeared less clearly defined compared to controls (arrowheads in Figure 5 and 6). Whereas the roof plate is normally devoid of differentiated MTN neurons, we observed MTN cell bodies within the roof plate following ROBO2 knockdown. This phenotype was not observed in GFP control electroporations (n = 11), indicating that it is a specific consequence of reduced ROBO2 expression.

Together, these findings demonstrate that SLIT-ROBO signaling is required not only for proper LLF axon pathfinding, but also for maintaining MTN neuronal somata adjacent to the roof plate and preventing their migration into it.

## Discussion

### Repulsion of MTN axons

During normal development, MTN neurons arise lateral to the roof plate, and their axons extend ventrally before turning caudally to pioneer the lateral longitudinal fascicle (LLF). MTN axons do not cross the roof plate into the contralateral side of the midbrain. This suggests that roof plate–derived signals prevent such misprojections. In the chick, SLIT proteins have been implicated in this process, both preventing inappropriate midline crossing and contributing to posteriorly directed growth toward the rhombencephalon (Molle et al., 2004; Farmer et al., 2008). Our co-cultures and transplantation experiments support this model, demonstrating that the roof plate is strongly repulsive for MTN axons, with SLIT2 contributing significantly to this activity.

Unexpectedly, knockdown of ROBO2 in MTN neurons did not lead to widespread axon crossing of the roof plate. Instead, axons displayed disorganized trajectories, with some misdirected toward the floor plate or even into the diencephalon, and only a small fraction crossing the midline. These findings suggest that while Slit2 – Robo2 signaling is critical for proper ventral and very likely posterior guidance, it is not solely responsible for preventing midline crossing. Additional roof plate – derived factors are probably there and act in concert with Slit2 to maintain ipsilateral axon trajectories.

In the spinal cord, Bmp7 has been shown to repel commissural axons (Augsburger et al., 1999), and BMP family member are expressed in the roof plate. However, our experiments do not support a major role for BMP signaling in MTN axon guidance. Application of BMP - containing beads did not alter MTN axons trajectory.and overexpression of BMP4 or GDF7 in dorsal midbrain failed to induce repulsion. These findings argue against BMP4 and GDF7 as major repellents for MTN neurons. Moschou et al., (Moschou et al., 2026) reported disruption of MTN axon growth in some cases after SMAD6 overexpression. SMAD6 is an intracellular inhibitor of BMP / TFG-ß signal transduction. Thus, the TGF-ß pathway might be involved in MTN axon growth direction.

In contrast, our *in vivo* transplantation experiments confirm that SLIT proteins exert a strong repulsive effect on MTN axons, consistent with previous studies (Molle et al., 2004; Kim et al., 2015; Ware, Schubert, 2011; Chedotal et al., 1995; Farmer et al., 2008). Both, Slit1 and Slit2 are expressed in ventral midbrain, diencephalon, and hindbrain and function as both short-and long-range repellents across species (Rajagopalan, Nicolas et al., 2000; Rajagopalan, Vivancos et al., 2000; Brose, Bland et al., 1999; Simpson, Kidd et al., 2000). In addition to preventing invasion of the floor plate, Slit signaling very likely contributes to restricting MTN axons from entering the diencephalon. Our *in vivo* transplantation experiments further support this, as both diencephalic and notochord tissue exerted repulsive effects on MTN axons. Interestingly, roof plate transplants had an even stronger effect, in some cases overriding diencephalic repulsion and redirecting axons anteriorly.

One possible explanation is that the roof plate transplants release morphogens such as BMPs, WNTS and FGFs, which may alter the surrounding tissue environment, potentially including the regulation of SLIT expression (Chizhikov et al., 2004). Because our transplants were performed at early developmental stages, prior to full dorsoventral regionalization of the midbrain (Li et al., 2005), they may have influenced local tissue specification. This raises the possibility that additional, as yet unidentified, repulsive cues are produced by or induced around the roof plate.

Another candidate repellent is the axon guidance molecule Draxin, which is expressed in a high - dorsal to low - ventral gradient in chick midbrain (Islam et al., 2009) and has been shown to repel MTN axons *in vitro* (Naser et al., 2009). Our unpublished data (Schmid et al., unpublished) suggest that *in vivo* overexpression of Draxin disrupts MTN axon trajectories and can lead to misrouting. This supports the idea that Draxin contributes to ventral guidance and may also help prevent MTN axons from crossing the roof plate. A similar inductive mechanism may underlie the effects of notochord transplants. The notochord is known to induce floor plate identity (Charrier et al., 2002; Van Straaten et al., 1985; Ruiz I Altaba et al., 1995), which produces different morphogens and axon guidance proteins (Colamarino & Tessier-Lavigne, 1995; Chedotal, 2019). The observation that MTN axons avoid not only the transplant itself but also the surrounding area suggests that secondary induction of repulsive signals, such as Slit proteins, may occur. Although Netrin1, an attractive cue, might be induced (Kennedy et al., 1994), its effects could be overridden by stronger repulsive signals. Indeed, Slit signaling has been shown to suppress Netrin - mediated attraction in other systems, such as during migration of inferior olivary neurons in the rhombencephalon (Causeret et al., 2002).

Taken together both, our co - culture and transplantation experiments indicate that MTN axons navigate within an environment rich in repulsive cues that constrain their trajectory. SLIT2-ROBO2 signaling plays a central role in this process, particularly maintain axon positioning relative to the roof plate. However, it likely acts in combination with additional factors, such as Draxin and possible other roof plate-derived signals, to ensure the MTN axons remain ipsilateral and correctly establish the LLF.

### ROBO deficiency changes MTN cell body location

Our experiments showed that the knockdown of ROBO2, a receptor for SLIT2, disrupts the normal ventral trajectory of MTN axons, causing them to lose their straight ventral orientation. Interestingly, ROBO2 knockdown also led to a striking change in neuronal positioning. MTN somata invaded the roof plate, thereby disrupting its typical neuron - free architecture. These findings indicate that, in chick midbrain, SLIT2–ROBO2 signaling is required not only for axon guidance but also for maintaining neuronal MTN somata in their correct lateral positions adjacent to the roof plate.

This result parallels findings from the mouse floor plate, where Robo2 prevents neuronal somata, including motor neurons, from invading the midline (Kim et al., 2014). Slit proteins have been implicated in regulating the migration of other neuronal populations in the developing brains. For example, in the rat olfactory system, Slit proteins produced by the septum repel subventricular precursor cells (Wu et al., 1999). In humans, Slit signaling regulates the migration of neural progenitors in the subventricular zone (Guerro-Cazares et al., 2017), and in mice, it controls midline crossing of precerebellar neurons in mice (Di Meglio et al., 2008; Dominici et al., 2018). These studies support a conserved role for Slit - Robo signaling in restricting neuronal migration relative to the midline.

Our finding also agree with the broader role of Slit – Robo signaling in longitudinal axon guidance. Robo1 and Robo2 receptors exhibit distinct but partially overlapping expression patterns: Robo1 is predominantly expressed in ventral and lateral tracts, whereas Robo2 is restricted to more lateral tracts in mouse (Kim et al., 2011; Long et al., 2004). Functional studies in mouse have shown that loss of Robo1 leads to increased axon occupancy in lateral tracts, while loss of Robo2 results in a shift of axons towards more ventral positions (Jaworski et al., 2001; Long et al., 2004). Combined loss of Robo1 and Robo2 causes severe axon guidance defects, including midline crossing and aberrant wandering (Farmer et al., 2008; Mastick et al., 2010; Sanhueza et al., 2025). Thus, Robo1 primarily regulates ventral tract positioning and midline exclusion, whereas Robo2 is critical for the guidance of more dorsal tracts such as the LLF (Farmer et al., 2008; Kim et al., 2011). Robo2 has also been shown to prevent ventral neuronal somata from invading the floor plate (Kim et al., 2011), a function that closely mirrors our observations in MTN neurons following ROBO2 knockdown around the roof plate.

In addition to their repulsive functions, Slit proteins have been reported to promote branching and elongation of sensory axons and cortical dendrites (Wang et al.,1999). Although we did not directly assess these effects, the disorganised axonal trajectories observed after ROBO2 knockdown may, in part, reflect altered responsiveness to such Slit-mediated growth-modulating signals.

Taken together, our findings show that Slit2 – Robo2 signaling plays a crucial role in directing the ventral growth of MTN axons in the chick midbrain. In addition, our findings identify SLIT2 as a key repulsive factor in maintaining the spatial segregation of MTN neuronal somata by preventing their invasion of the roof plate. Thus, Slit2 expression in the roof plate is essential not only for axon pathfinding but also for preserving the proper organization of MTN neuronal somata relative to the midline.

## Acknowledgements

We are grateful to Lothar Just for helpful disussions and comments on the manuscript. Dr. H. Lieckert for the figt of the VenusLyn reporter plasmid and Dr. Avihu Klar for the plasmid containing a ggROBO sequence.

## Author contribution

Conceptualization: A.W., A.G.; Investigation: C.L., A.K., M.N., S.R., U.K.; Writing: C.L., A. K., M.N., A.G., A. W.; Funding acquisition: A.W. B.H.

## Animal ethics

This study was conducted in strict accordance with the Federal Republic of Germany (TierSchG) and the Animal Care and Use Committee (ATV) of the local authorities (Regierungspräsidium Tübingen). Ethical approval was not required for this study in accordance with local regulations governing the use of early-stage avian embryos. Notwithstanding, the chick embryos were treated with respect. All efforts were made to minimize any suffering.

